# From topography to connectome: Towards an integrated understanding of the resting brain

**DOI:** 10.64898/2026.06.06.730493

**Authors:** Samuel Naranjo Rincón, Fyzeen Ahmad, Ty Easley, Shirin Shoushtari, Tristan Glatard, Gregory Kiar, Hailey Modi, Simon Dahan, Emma Robinson, Ulugbek S. Kamilov, Janine Bijsterbosch

**Affiliations:** Department of Radiology, Washington University Saint Louis, MO; University of Minnesota, Minneapolis, MN; Krembil Centre for Neuroinformatics, Centre for Addiction and Mental Health, Toronto, Canada; Center for Data Analytics, Innovation, and Rigor, Child Mind Institute, NY; Research Department of Biomedical Computing, School of Engineering & Imaging Sciences, King’s College London, UK; Centre for Neurodevelopmental Biology, Institute of Psychiatry, Psychology & Neuroscience, King's College London, UK; Department of Electrical & Systems Engineering, Washington University Saint Louis, MO; Department of Electrical and Computer Engineering, University of Wisconsin-Madison, WI

## Abstract

As the field expands from early research into the human connectome, there has been a fast expansion in the number of analytical approaches to study resting state functional MRI (rsfMRI) data. With increasing focus on individual differences, topographical brain maps of spatial organization have emerged in addition to traditional functional connectomes. Here, we developed a deep-learning model to embed maps of network topography and faithfully translate to individualized connectomes. Results confirmed the validity of the surface vision transformer based on reconstruction accuracy (0.73±0.09) and accurate topography-to-connectome translation (0.43±0.08). Importantly, translated connectomes retained identifiability and brain-cognition associations. These findings establish a direct mapping from spatial topography to connectomes that can be used to integrate scientific insights across rsfMRI sub-fields. This is an important step towards broadening our conceptualization of the connectome and supporting a broader integration of findings to inform a complete understanding of the human connectome.

**Teaser:** Translating from spatial maps of brain organization to region-to-region connectomes retains shared individual differences.

## Introduction

The human connectome was initially proposed as a way to illustrate how functional brain states emerge from their structural substrates (*1*). Specifically, the goal of the human connectome was to provide a robust structure-function mapping of the human brain by combining information about white matter structural connectivity (SC) from diffusion weighted imaging with functional connectivity (FC) from resting state functional magnetic resonance imaging (rsfMRI). Over the last 20 years, the human connectome has helped characterize various properties of SC such as network organization (*2, 3*), its role in clinical settings (*4*), and developmental changes (*5*–*7*). For FC, this structure-function mapping revealed large overlaps of pathways with SC across the brain (*8*–*10*). However, not all FC connections are bound by known cortico-cortical pathways, largely due to FC representing statistical co-activation patterns across the cortex (*9, 11*). Consequently, the connectome has offered a flexible definition of FC, contributing to its rising popularity in brain-wide association studies, and resulting in a rapid expansion of approaches to study FC.

Generally, FC is studied using group-derived parcellations to extract time series and calculate region-by-region temporal correlations (*12*). Beyond these functional connectomes, studies have expanded the quantification of rsfMRI to derive individualized spatial topography maps (*13*). This expansion of possible brain representation (i.e., low-dimensional summary features) derived from rsfMRI scans has provided novel insights into individual differences in functional organization (*14*). However, this analytical variability also catalyzed divergent methodological silos, limiting replicability and clinical applications (*15*). Direct comparisons of results have proven helpful to map out the effects of analytical variability in related domains including fMRI preprocessing and relevance to behavior (*14, 16*). However, leveraging direct comparisons to integrate FC findings is challenging due to fundamental mismatches in the type of brain representations (temporal connectomes vs topography maps), as well as their dimensionality and spatial boundaries. Despite these fundamental mismatches that obscure straightforward comparison, both brain representations are derived from the same rsfMRI scans and may therefore capture overlapping information.

There is an important need to integrate findings regarding functional organization across different rsfMRI brain representations, such as region-to-region temporal connectomes and topography maps. Such integration of rsfMRI findings is necessary to guide correct mechanistic interpretations, gain a comprehensive understanding of the connectome, and inform best practices for analytical approaches in clinical trials. Recently, a toolbox was developed to assess spatial correspondence between group-level parcellations used to define temporal connectomes and group-level topography maps (*17*). These results are useful for comparing group-level spatial parcellation definitions, potentially in combination with standardized nomenclature (*18*). However, this spatial correspondence toolbox was not designed to integrate findings regarding individual differences. Other studies have anchored comparisons of individual differences across rsfMRI brain representations based on their associations with behavior (*19*–*21*). These studies showed remarkable consistency in the behaviorally-relevant information captured regardless of the parcellation and indicated 60% shared variance between temporal connectomes and topography maps (*21*). Although these results are valuable to map behaviorally relevant shared variance between brain representations, brain wide associations have been shown to capture a relatively minor portion of individual differences, such that the use of behavior to anchor comparisons has its own limitations (*22*).

In our prior work, we leveraged topological data analysis to identify how brain representations compare in preserving inter-subject variability features on a shared manifold (*23*), revealing overlap between topography and amplitudes but dissimilarity between topography and partial FC connectomes. Others have opted to facilitate comparisons across brain representations by developing transforms. For example, leveraging the optimal transport problem, parcellated time series can be directly transformed from one atlas into another (*24*). Furthermore, Krakencoder has successfully translated across functional and structural connectomes, offering a promising pathway to facilitate cross-pollination of connectome-based results (*25*). However, current translation efforts have only been applied to parcellated timeseries or connectomes (*24*–*26*). Unlike connectomes, topography maps (such as maps derived using dual regression after group Independent Component Analysis; ICA) are represented as one high-dimensional cortical sheet for each map, typically ranging from 15 to 50 topography maps (*27, 28*). These differences render current rsfMRI translations unsuitable for topography to connectome translations. Here, we expand beyond atlas or connectome transforms to develop a deep learning model that translates individual-specific spatial topography maps into respective individualized connectomes.

We introduce a deep learning model capable of embedding maps of network topography into a latent low-dimensional space from which the model can faithfully translate to an individualized connectome. Our encoder-decoder deep learning architecture (Fig. 1) builds on prior work developing the Surface Vision Transformer (SiT) that applies multi-head attention (*29*) across all vertices on any non-genus surface (*30*). Briefly, our findings validate the SiT as an appropriate encoder framework when embedding cortical surface maps, allowing us to achieve topography-to-connectome translation accuracy on par with prior connectome-to-connectome translations while preserving downstream brain wide association results. These findings therefore establish an efficient direct mapping from spatial topography to connectomes of rsfMRI, providing a path to integrate insights and results across the two.

**Fig. 1.**
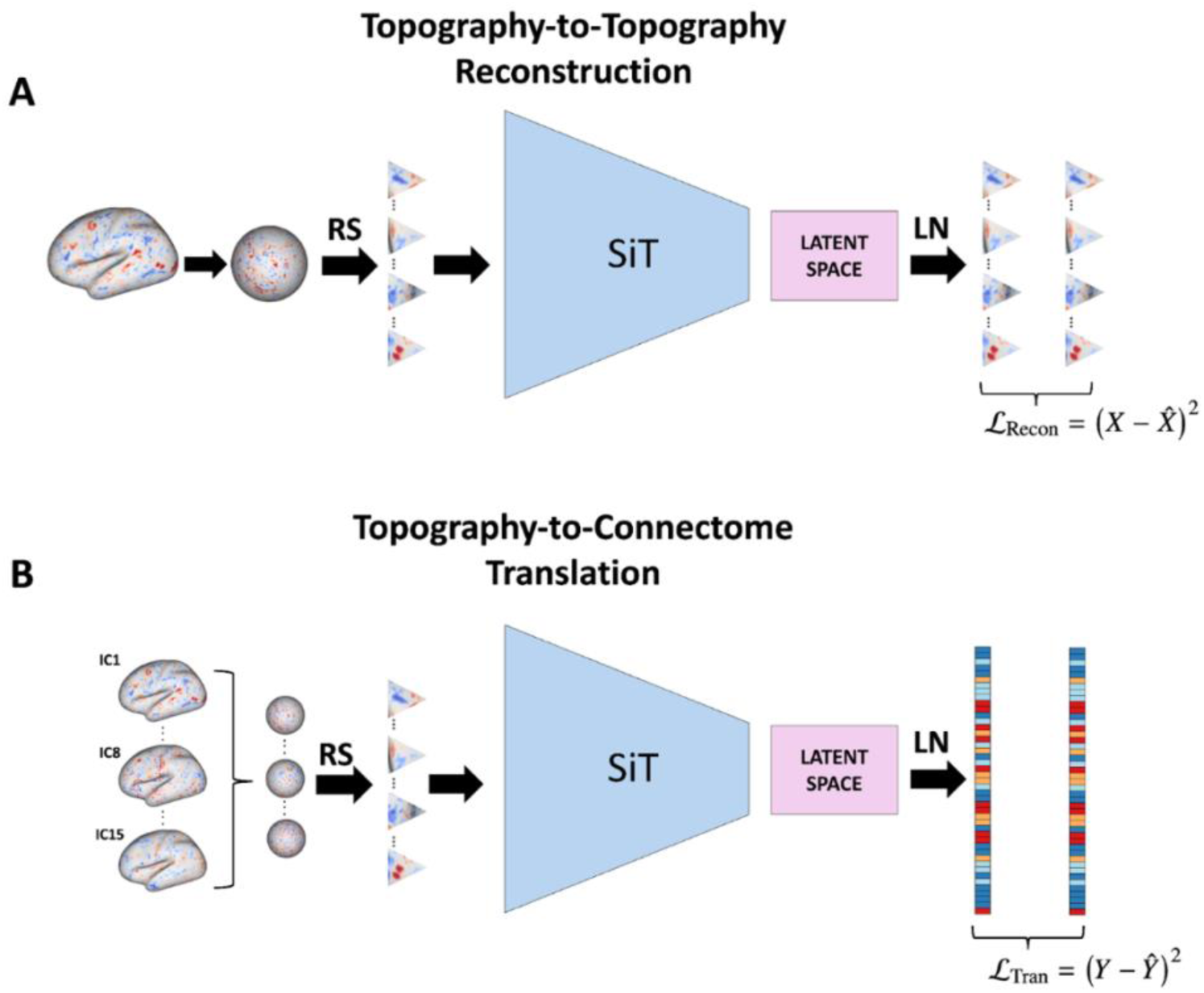
Schematic for ICA embeddings into latent space and linear decoders. (**A**) Topography-to-topography models embedded the vertex space of each subject’s normalized topography map (separate models for each of 15 ICA networks) into a lower dimensional triangular tessellation that formed the input to the SiT. Output of the SiT was mapped into a latent vector from which a linear transformation reconstructs triangular patches that were compared to the measured normalized topography input patches. (**B**) The translation models were trained using the normalized measured topography maps of all 15 ICA networks as input. Each topography map was first resampled into the same non-overlapping triangular tessellation patches used in (A), which were combined across all 15 spatial brain maps. The models then projected these patches into a latent space and used a final linear transform to decode the upper triangle of the demeaned FC connectome, which was compared to the measured demeaned FC connectome. Importantly, the group mean was removed from the inputs to the models and the targets for reconstruction or translation to avoid regression to the mean and encourage sensitivity to individual differences. Abbreviations: Reshape (RS), Surface Vision Transformer (SiT), Linear Layer (LN).

## Results

We trained and tested our rsfMRI topography-to-connectome translation framework (Fig. 1) using data from the Adolescent Brain Cognitive Development (ABCD) cohort with a site-based split of training (N_train_ = 7104), validation (N_val_ = 920), and test (N_test_ = 624). Model performance was primarily assessed using the within-subject correlation between measured and translated outcomes after subtracting the training group mean.

The result section consists of four main parts. In the first part, we validated the SiT encoder architecture by assessing reconstruction accuracy in a topography-to-topography framework for individual-specific topography maps. In the second part, we applied the validated encoder into a topography-to-connectome framework to translate from a 15-dimensional set of ICA topography maps to a 100-dimensional connectome. Our goal was to translate these individual-specific topography maps into corresponding individualized connectomes. In the third part, we therefore explicitly tested the preservation of individual differences by assessing to what degree the translated connectomes retained subject identifiability and brain-cognition associations. A translation model is only practically useful if it translates well to different settings. Therefore, in the fourth part, we determined the generalizability of the topography-to-connectome model to different connectome dimensionalities (100 vs 300), and parcellations (Schaefer vs Glasser). Full details on the architecture, hyperparameters, optimization, and assessments are available in the Materials and Methods.

### Encoder validation based on reconstruction of spatial topography

We began by validating the SiT as an encoder for brain topography. To assess performance, we trained 15 different encoders to separately reconstruct each of the 15 ICA individual-specific brain maps (Materials and Methods). Importantly, the reconstruction target topography maps were normalized to avoid regression to the mean and encourage accurate reconstruction of individual differences. Reconstruction accuracy was assessed as the bivariate correlation between normalized measured and reconstructed maps.

Across subjects, SiT encoders achieved a high reconstruction accuracy of r = 0.73±0.09 across all components (Fig. 2A). While individual differences were successfully reconstructed across brain maps, there was some variability in reconstruction performance. Of the 15 brain maps, brain map 1 (visual network; supplementary Fig. S1) had the highest reconstruction performance and the lowest intersubject variability (r = 0.88±0.01). Importantly, the SiT was able to perform well in other more heterogeneous networks such as the Default Mode Network (DMN; brain map 4; supplementary Fig. S1), with a reconstruction accuracy of r = 0.76±0.2. Example subjects for reconstruction in the top 10%, median, and bottom 10% are shown in Fig. 2. High reconstruction accuracy of individualized spatial topography maps shows that the SiT encoder was successful at embedding each brain map into a lower dimensional latent space. Moreover, these embeddings were robust to varying degrees individual differences, such that both homogenous primary regions and more heterogenous associative regions preserved a high degree of the original spatial organization. In the next section, we leveraged the SiT encoder to translate from spatial topography to connectome.

**Fig. 2.**
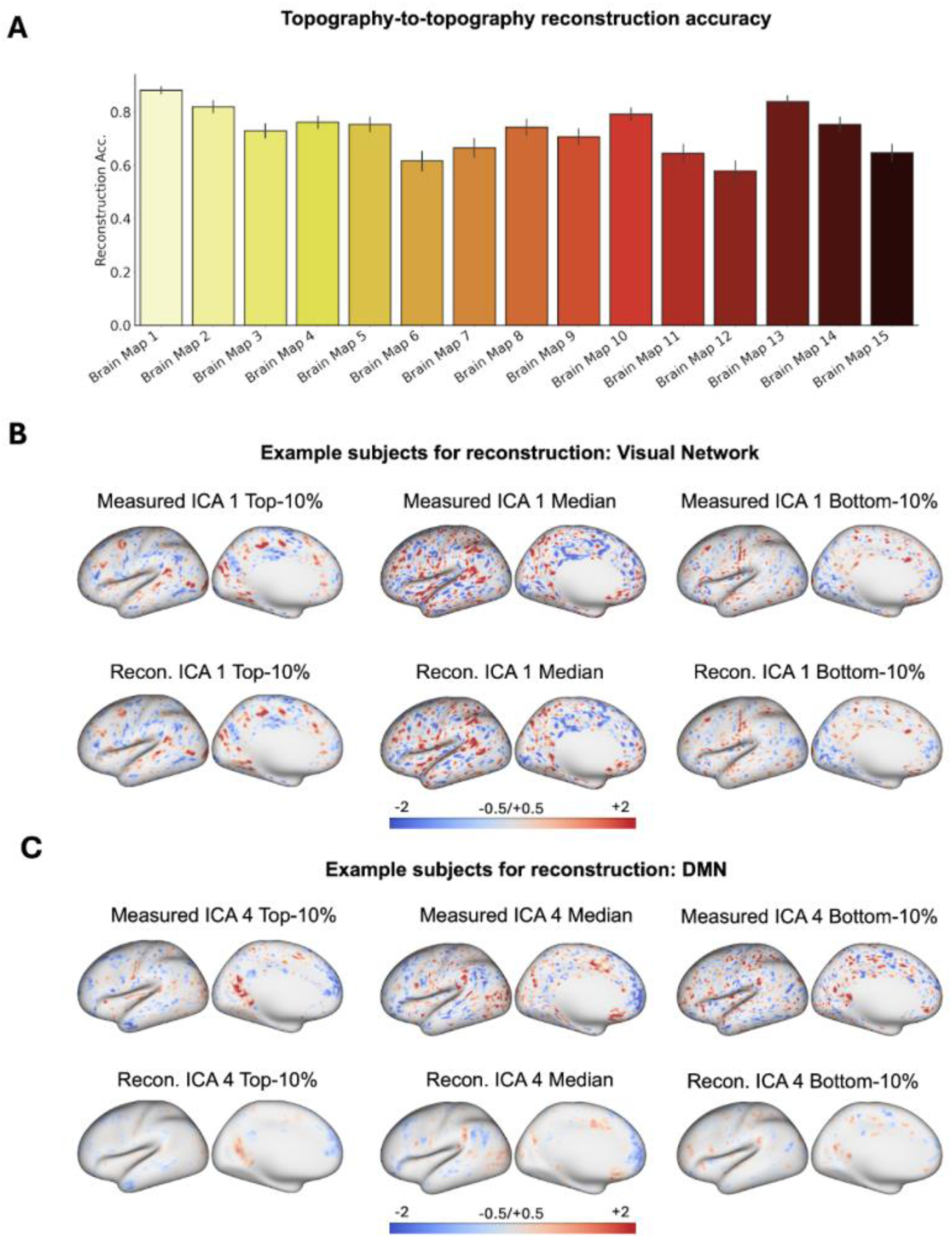
Topography-to-topography reconstruction accuracy and example subjects. (**A**) Each of the 15 topography maps achieved high reconstruction accuracy. See supplementary Fig. 1 for group ICA maps corresponding to the 15 brain maps. The barplot shows mean across subjects with error bars showing one standard deviation. (**B**) Three examples of normalized topography maps (measured and reconstructed) for selected example subjects in the Top-10%, Median, and Bottom-10% accuracy range for brain map 1. Brain map 1 corresponds to the Visual Network, and individualized maps are normalized to focus on individual differences. (**C**) Three examples of normalized topography maps (measured) for different selected subjects in the Top-10%, Median, and Bottom-10% for brain map 4, corresponding to the Default Mode Network. Measured maps were normalized by subtracting the training data mean (and division by the training standard deviation). These normalized maps were used in the model as inputs to focus accuracy on individual differences.

### Translating from spatial topography to connectomes

We combined the SiT encoder with a linear decoder to translate from topography network maps (all 15 ICA maps) to a connectome defined as the full pairwise bivariate correlations using the 100-dimensional Schaefer atlas (*31*) (Materials and Methods). Similar to the reconstruction application, we trained the model to translate demeaned connectomes to avoid regression to the group mean. Translation accuracy was assessed with the demeaned versions of both the measured upper triangle of the connectome and its translated counterpart.

Translation accuracy in the held-out test dataset was on average r = 0.43±0.08. Connectome translations for representative subjects in the top 10%, median, and bottom 10% are shown in Fig. 3. Other metrics of model performance are shown in supplementary Table 1, and model performance as a function of training mini-batches in training and validation data is shown in supplementary Fig. S2. Taken together, these results confirm the ability of the SiT encoder-decoder framework to translate from spatial FC topography to individualized connectomes. In the next section, we explicitly evaluated to what degree individual differences were retained after translation.

**Fig. 3.**
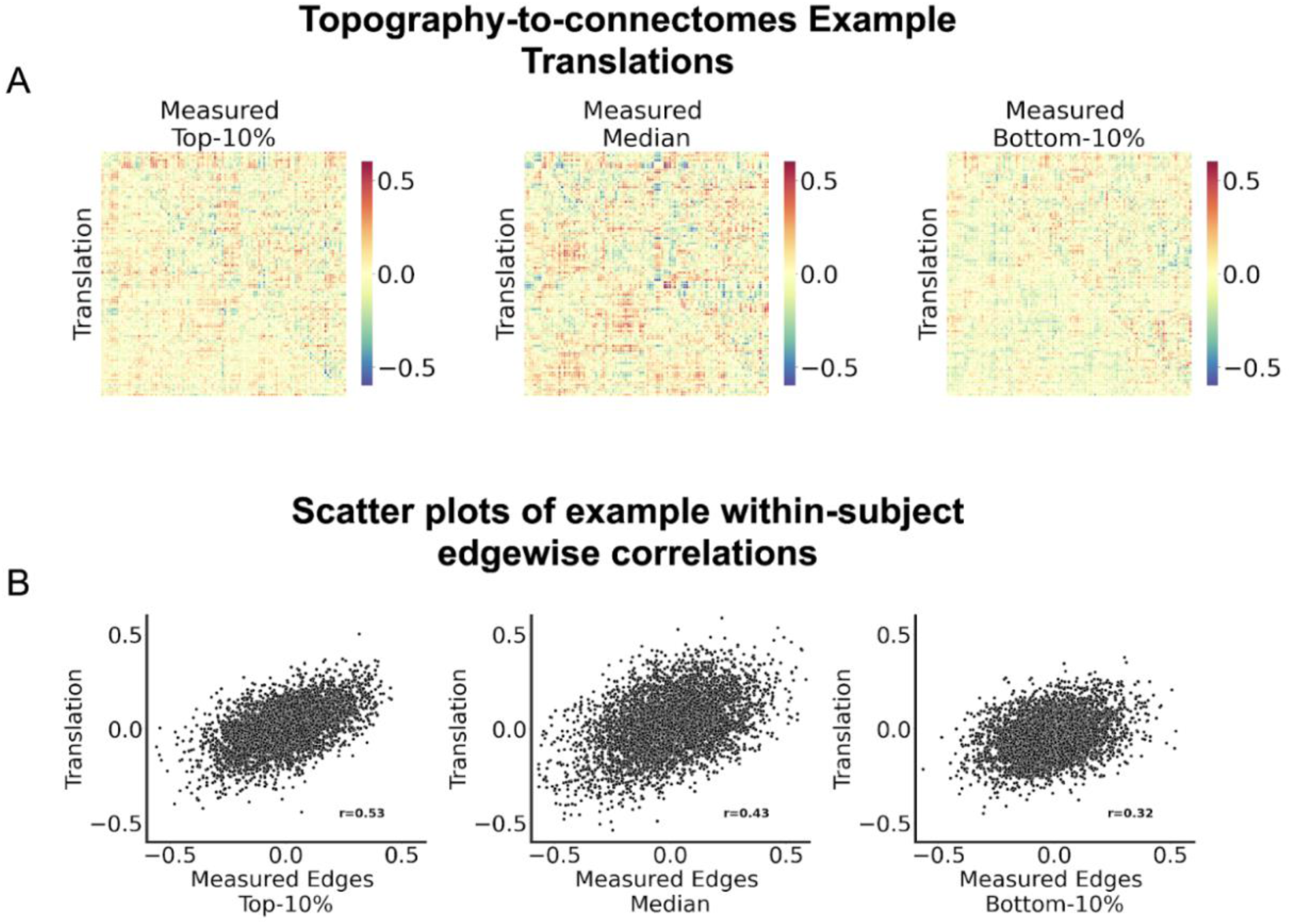
Translation accuracy for example subjects. (**A**) Three examples of demeaned connectomes for selected example subjects in the Top-10%, Median, and Bottom-10% accuracy range. Demeaned connectomes are shown to focus on individual differences. Translated connectomes are the lower triangle and measured connectomes populate the upper triangle. (**B**) Scatter plots of same connectome edges, showing degree of overlap between measured and translated connectomes for those same example subjects. Correlation between measured and predicted edges (i.e., the within-subject translation accuracy) is shown at the bottom right. Left: Random top 10% subject example. Middle: Median subject example. Right: Random bottom 10% subject example.

### Preserving connectome individual differences

A key goal of our translation architecture was to retain individual-specific features of subject connectomes. As such, we tested the preservation of individual differences across translated connectomes. Specifically, we performed two different tests of subject identifiability and two separate brain-cognition association analyses. Firstly, we assessed the average rank percentile, which compares how a measured connectome and its corresponding translation compare against all other translated connectomes. To successfully preserve connectome individual differences, any given measured connectome should be more correlated with its translated counterpart than with as many other translations as possible. Average rank percentile measures this fraction, where a ratio of 1 reflects perfect subject identifiability based on translations only. Secondly, we tested the Top-1 accuracy, which is a binarized measure per subject where a positive outcome requires that a given translated connectome has a higher similarity score with that individual’s measured connectome than with all other translated connectomes. As such, the Top-1 accuracy represents the proportion of subjects with a perfect rank percentile of 1 - reflecting a higher bar than the average rank percentile. Thirdly, we used the NIH toolbox composite scores for crystallized and fluid intelligence to assess whether brain-cognition associations in translated connectomes matched measured connectomes. We performed univariate edgewise brain-cognition associations for both measured and translated connectomes separately and quantified their similarity with the bivariate correlation coefficient. Lastly, multivariate brain-cognition analyses were performed using principal component regression (PCR) to predict cognitive scores. These multivariate results were repeated 1000 times using a 90/10 train-test shuffle split within the test (N_test_ = 624) dataset to assess if translated connectomes preserved measured multivariate brain-cognition associations.

Average rank percentile across subjects showed near perfect subject identifiability (mean = 0.99; Fig. 4A). Most subjects had their translated connectomes match their measured connectomes more strongly than any other subject, and those that did not achieve 100% identification still remained at a range of 0.96 - 0.99 (Mean = 0.99), illustrating the topography-to-connectome model’s ability to preserve individual differences for finger printing. These results were echoed by identifiability results based on Top-1 accuracy for the translated models. Specifically, for 53% of participants their translated connectomes were more highly correlated with their respective measured connectome than with any other translated connectome, compared to a 0.16% chance level (1/624).

**Fig. 4.**
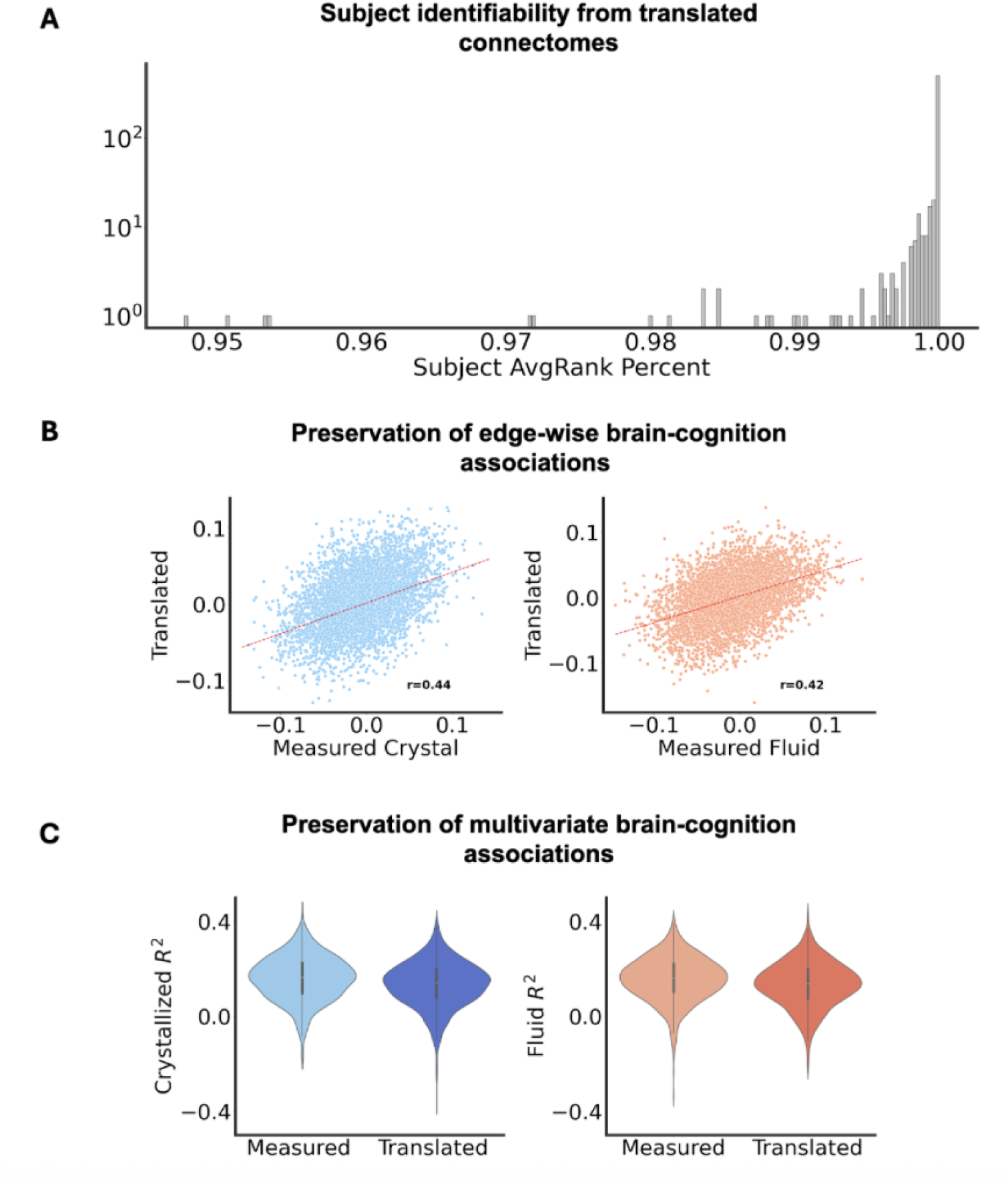
Preserving individual differences in translated connectomes. **(A)** Average-rank percentiles for Schaefer-100 translated connectomes reveal high identifiability (average rank = 0.99). Here, the figure shows a histogram of subject counts on a logarithmic y-axis scale. **(B)** Univariate brain-cognition results capture similar individual differences. Scatter plots show the distribution of edge-wise correlations with cognition for measured and translated connectomes for crystallized intelligence (left) and fluid intelligence (right). Red dotted lines are a fitted 1D polynomial regression minimizing the least squares error, showing positive correlation between measured and translated results. Bivariate-correlation coefficients are shown on the bottom right. Multivariate results show highly similar prediction accuracy when using measured connectomes (left violins, light shade) compared to translated connectomes (right violins, dark shade) for both crystallized intelligence (left-hand plot) and fluid intelligence (right-hand plot). Violin plots show R2 scores distributed over 1000 shuffle splits. Inside violin plots are box plots showing the mean and inner quartiles of scores.

Univariate comparisons revealed preserved brain-cognition associations between measured and translated connectomes for both crystallized (r = 0.44; Fig. 4B) and fluid (r = 0.42; Fig. 4B) intelligence. Multivariate analyses for translated connectomes revealed an average cross validated accuracy of r^2^ = 0.16±0.1 (Measured: r^2^ = 0.17±0.1; Fig. 4C) for crystallized intelligence and r^2^ = 0.11±0.09 (Measured: r^2^ = 0.15±0.09; Fig. 4C) for fluid intelligence when using translated connectomes. These results show that translated connectomes sustained sufficiently robust signals to preserve similar brain-cognition associations unique to individual subjects’ connectome profiles.

### Generalizability of topography-to-connectome translations

We performed multiple analyses to assess replicability. Here, we trained three new models (Fig. 1B) to predict different connectomes from the 15 ICA topography maps. Specifically, we assessed the topography-to-connectome framework’s ability to translate to higher-dimensional connectomes (Schaefer-300 and Glasser-360) and partial FC connectomes.

The results of re-trained models for different connectomes revealed a slight drop in performance when translating higher-dimensional connectomes. Specifically, the Schaefer-300 translation accuracy was r = 0.35±0.06 (Fig. 5A) and the Glasser-360 translation accuracy was r = 0.32±0.06 (compared to r = 0.43±0.08 for Schaefer-100; Fig. 5A). Notably, the translation accuracy was substantially lower for partial FC connectomes (r = 0.15±0.04; Fig. 5A). Overall, translated connectomes, across different dimensions and FC types, had strong average rank percentile identifiability (Fig. 5B). Top-1 Accuracy was harder to achieve, but most models performed above chance (12% Schaefer-300; 24% Glasser-360; 0.16% Chance; Fig. 5C). Translated Schaefer-100 partial FC connectomes had the lowest Top-1 accuracy score of 0.04%.

**Fig. 5.**
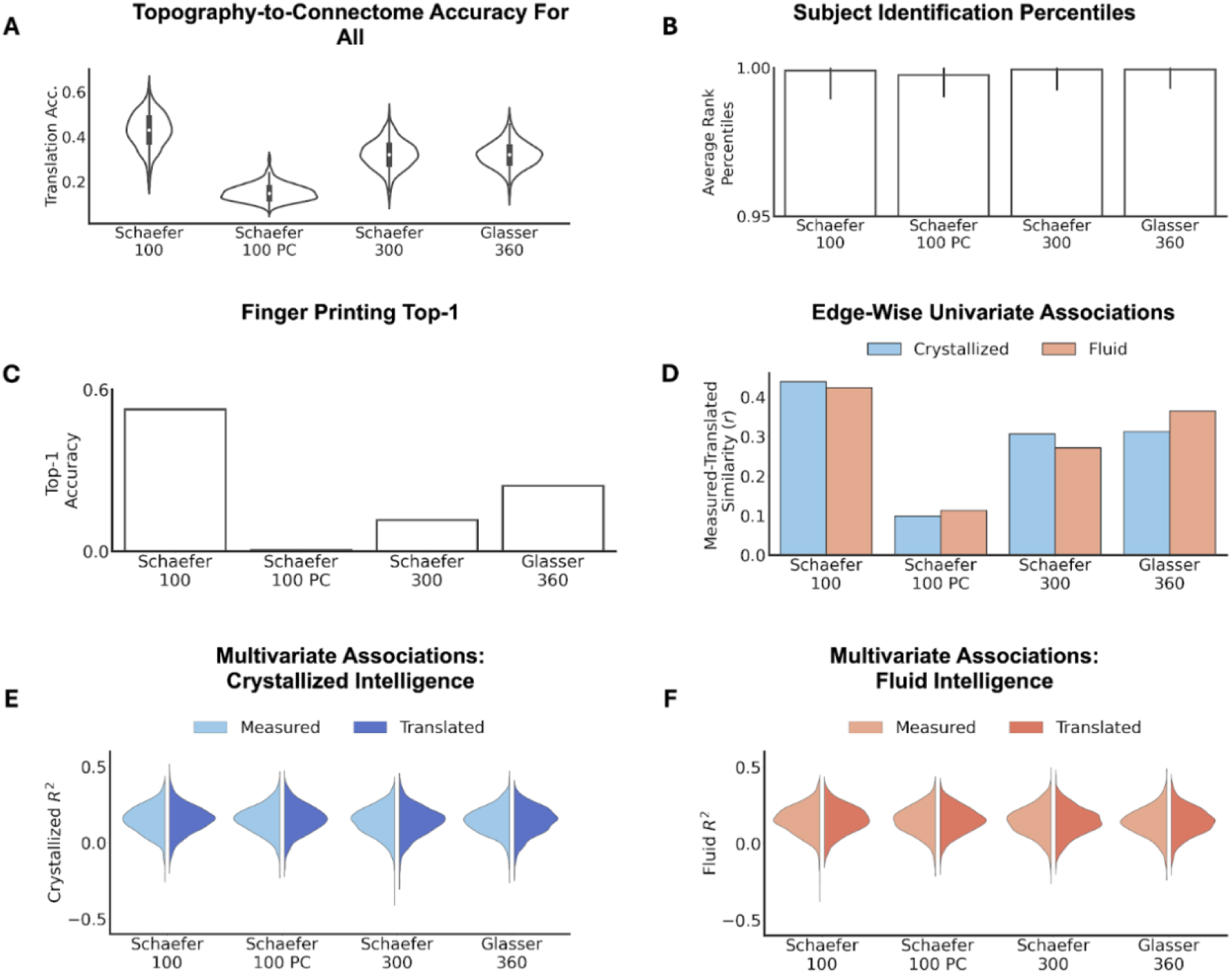
Model performance when translating to other connectome types. Retraining the model to translate different connectomes from the same topography maps showed mixed results. (A) Translation accuracy was largely retained for higher dimensional connectomes and across atlases, but was lower when translation partial connectomes (PC). (B) Average rank percentile identifiability was high for all translated connectomes. Here, good performance for partial connectomes (despite poor accuracy; A) suggests that the added noise from poor translation does not make a subject more similar to other subjects. Bar plots show the mean across subjects and error bars reflect one standard deviation. (C) Top-1 accuracy dropped for higher dimensional connectomes and was close to zero for partial connectomes (PC), in line with accuracy results (A). Univariate brain-behavior associations follow a similar pattern with lower edge-wise correlations between measured and predicted associations. (E) Multivariate brain-cognition associations show similar explained variance between measured (left violins, light shade) and translated (right violins, dark shade) connectomes for crystallized intelligence. (F) Multivariate results for fluid intelligence were similar to crystallized intelligence (E). Violin plots in E and F show results across 1000 shuffle split cross validations.

Brain-cognition associations across both connectomes were similar but bounded by their translation accuracy. Univariate analyses on translated Schaefer-300 and Glasser-360 connectomes showed moderate similarity with analogous measured edge-wise univariate associations for both crystallized and fluid intelligence (Fig. 5D). Translated partial FC connectomes again showed the lowest similarity between measured and translated univariate results. Multivariate analyses revealed similarity between translated and measured connectomes for both cognitive measures (Fig. 5E). Our topography-to-connectome translation architecture achieved faithful translations of measured connectomes, not only edge-wise values of individualized connectomes, but also downstream analyses and results. This performance extended across atlas dimensions and atlas type.

## Discussion

In this work, we introduced a novel topography-to-connectome translator model that can map from individualized spatial maps to analogous connectomes, while preserving individual differences in finger printing and downstream brain-cognition analyses. We first validated the proposed spatial topography encoder, showing high and consistent reconstruction accuracy across brain maps, including both primary and higher association networks. When testing the full topography-to-connectome translator, translated connectomes demonstrated consistent and robust preservation of measured connectomes including preserved subject identification and brain-cognition associations. Overall, this paper contributes to the connectomics field by integrating individual difference findings from spatial topography analyses of rsfMRI data with findings from FC connectomics.

Our topography-to-connectome translator achieved a performance of r = 0.43 (demeaned correlation between measured and translated connectomes), well within the range of previous connectome-based translation efforts. For example, Krakencoder was optimized to map from one connectome type to another, using a many-to-many approach by embedding various kinds of connectome “flavors” and assessing reconstruction and translation accuracy among them (*25*). In their study, the accuracy for reconstructions and translations ranged r = 0.30-0.70, except for partial FC connectomes (typically r = 0.07-0.30). Others have attempted to translate across SC and FC by using a more complex generative model. For example, authors in (*26*) trained an adversarial generative model to translate from structural and functional connectomes, with performance accuracies typically ranging from r = 0.34-0.43. Using our SiT encoder-decoder, translation accuracy somewhat diminished when translating to higher resolution connectomes but was stable between different atlas types (Fig. 5). The small decrease in translation accuracy for the Schaefer-300 and Glasser-360 parcellations may result from having the linear decoder infer more elements for higher parcellation sizes. Additionally, translations to higher dimensional connectomes may be impacted by residual between-subject misalignment. Future work may wish to explore how improved alignment using multimodal surface matching (*32, 33*) prior to the calculation of topography maps may impact translation accuracy.

Our topography-to-connectome direction presents a different task from the connectome-to-connectome translations discussed above because the source is a non-euclidean object being mapped onto a Positive Semi-Definite connectome form. In fact, our previous work suggested that spatial maps and connectomes have predominantly diverging inter-subject variability features, making them difficult to compare (*23*). Despite this diverging variability, our translator converged to a non-linear mapping that directly translated from post-preprocessed dual regressed ICA brain topography maps onto post-preprocessed post-parcellated FC connectomes. As such, our trained model offers an accessible opportunity for a multiverse approach without the need for reanalysis (*34*). More broadly, the fact that we can approximate connectomes directly from topography (without timeseries) may illuminate important shared information between divergent analytical brain representations of rsfMRI data that could be leveraged to enrich and integrate our understanding of the resting brain. Connectomes define a binary assignment scheme where all vertices within a parcel are averaged to define the time-series for that parcel. The spatial definitions of parcels are often defined from a group-average atlas, whereas topography maps like ICA, define varying functional boundaries across the cortex and across subjects. Therefore, for any given group-averaged parcel, differing functional boundaries across subjects systematically impact the parcel time series and thus establish an association between parcel timeseries and functional boundaries in ICA that can be exploited by our topography-to-connectome translators (*35*).

Importantly, the translated connectomes retained individual difference information. Identifiability was strong, suggesting the usability of translated connectomes for finger printing (*36*). In fact, both univariate and multivariate brain-cognition associations preserved enough information regarding individual differences, to the extent that both measured and translated connectomes were strongly correlated in their univariate associations and achieved similar multivariate brain-cognition associations. These results suggest that connectome-wide individual differences were retained despite a slight drop in edge-wise individual differences. This advantage of multivariate over univariate performance may be driven by relatively higher levels of noise at the edge level and/or shared variance (e.g., redundancy of information) across the full multivariate connectome (*37*).

Notably, translating to partial connectomes had the lowest performance of all connectome translations. This result is somewhat surprising given that partial connectomes have been shown to benefit from noise reduction (*38*). However, as mentioned previously, similar drops in performance were observed in the Krakencoder (*25*), and partial connectomes have been shown to have lower test-retest reliability (*39*), potentially due to instability in regularization (*40*). Furthermore, ICA derived topography maps represent whole brain connectivity for any given network by grouping similar vertices, which more closely resemble full correlation connectomes. Partial correlation FC connectomes, instead, regress out all other time series before calculating a pairwise edge, resulting in a sparser FC matrix. This difference may explain why translation accuracy was higher for full correlation than partial correlation connectomes. Further work is needed to integrate partial connectome results with ICA and full correlation FC connectomics.

Integrating results across rsfMRI studies is crucial to inform translational research of brain diseases. To date, practical applications of rsfMRI in clinical settings are relatively limited. This dissonance between rsfMRI analyses and clinical work arises from limitations in brain wide association (*22*), non-standardized nomenclature (*18*), disagreement across analyses pipelines (*16*), and flexibility in choice of brain representation (*15*). The topography-to-connectome translator model addresses the latter issue by assuming brain representations share underlying latent features that can be used to tether them. Recently, this perspective has been expanded on to enable a “multiverse approach” without the high computational burden (*34*), by embedding various kinds of analysis pipelines into a lower dimensional latent space and using a deep learning approach to sample and learn across this subspace. Learning inter-relationships between pipelines from this low-dimensional space allowed authors to approximate a wide spectrum of rsfMRI analytical flexibility. Similarly, Krakencoder exploited the latent low-dimensions space of inter-relationships across connectome “flavors” (*25*). We expanded on this concept by converging to a low-dimensional latent space that merged topographies and connectomes for translations.

Although our results revealed the ability to translate from topography and connectomes while retaining individual differences, there are several limitations to discuss. Firstly, deep learning models require optimization of a large number of parameters, which requires a big training dataset. Although we achieved reasonable performance using ABCD data, it is possible that we have not yet hit the ceiling of translation accuracy. In future work, we plan to expand to the UK Biobank cohort (which now includes N=100,000, enabling a more than 10-fold increase in our current training sample) to test this hypothesis. Secondly, there are many other types of topography brain maps (additional dimensions and methods), many other connectomes, and additional sources of analytical variability (e.g., variation resulting from data acquisition and preprocessing pipelines). Although we tested the ability to translate to different connections, future work will be needed to expand the translations to a broader set of topography and connectome brain representations. Specifically, in future work, we plan to expand into a comprehensive set of ‘NeuroTranslators’ that separately learn mappings from one topography based representation to some connectome profile, such that each separate model can be fused to reach an abstracted unifying latent space capturing underlying statistical properties of rsfMRI data that facilitate direct translation and comparison.

In summary, we demonstrated the ability to faithfully translate individual FC connectomes from rsfMRI topography maps. These results broaden our conceptualization of the connectome and emphasize the relevance of rsfMRI brain representations that are not traditionally considered as ‘connectomes’ to the field of connectomics. As such, we believe a broader integration of findings will be valuable to inform a complete understanding of the human connectome, not only within the context of other types of connectomes, but beyond that and onto other domains of analyses. With this in mind, our topography-to-connectome mapping seeks to resolve a goal identified in the original proposal of the human connectome, namely the creation of a publicly accessible standardized data format. In the presence of expanding analytical approaches to study rsfMRI, this standardized data format goal can be reenvisioned through comparison and integration facilitated by translation.

## Materials and Methods

### Dataset

The Adolescent Brain Cognitive Development (ABCD) cohort is a publicly available dataset with a large sample size of typically developing children (N=8,648) from ages 9 - 11 recruited across 21 different sites in the United States (*41*–*43*). Data were acquired across three types of 3T scanners (Siemens Prisma, General Electric 750, and Philips) running standard echo-planar imaging sequence protocol (TR=800ms, TE=30ms, flip angle=52°, multiband factor=6, 2.4mm isotropic voxel resolution). ABCD data were preprocessed using the ABCD-BIDS pipelines (*42*), including standard frame-to-frame head motion correction, nonlinear alignment to standard space, Freesurfer extraction of the cortical ribbon, and formatting to grayordinate format. Data cleanup for rsfMRI furthermore followed the ‘DCAN-BOLD Preproc’ module, including regression of white matter, CSF, mean CIFTI timeseries, the six motion parameters and their volterra expansion, bandpass filtering (0.008-0.09 Hz), and motion censoring. Each subject had four 5-minute resting state scans, which when concatenated resulted in a total of 20 minutes of resting state data (*41*). To account for site and family effects, 14 sites were used as the training data, 3 as the validation data, and 3 as the testing data, resulting in a final split of N_train_ = 7104, N_val_ = 920, and N_test_ = 624.

### Brain Representations

Group ICA was performed on the rsfMRI data concatenated across all subjects at a dimensionality of 15, followed by dual regression to estimate subject-specific surface network maps (*27, 28*). The dimensionality of 15 was chosen to balance the topographical information available to models while maintaining computational feasibility of the SiT embedding. Additionally, to ensure only individual differences were embedded, each spatial brain map was normalized by the training sample mean and standard deviation for that brain map. For connectomes, each subject’s data was spatially parcellated into 100/300 regions based on the Schaefer atlas (*31*). The same process was done for the Glasser 360 atlas (*44*). Time series from each region were extracted and correlated with every other to construct a pairwise bivariate full correlation matrix. Partial correlations were produced by applying the Moore-Penrose pseudo-inverse on the full correlation matrix, which is analogous to using an L2-regularized inverse.

### Embedding and Translating Spatial Topography Maps

Both reconstruction and translation used the same SiT-based encoder paired with a linear decoder. We did not use a convolutional encoder because these have been shown to perform less well for cortical surface data (*30, 45*). The reason that transformers outperform convolutional neuronal networks (CNNs) can be traced to their powerful self-attention learning mechanism (*29, 46*) that encodes understanding of long-range context via modelling the co-variance of different features across the entire surface. This is necessary for modelling cognition since these processes recruit distributed networks spread over the entire cortical surface. CNNs cannot achieve this because they purposely learn highly localized filters. More recently, designs of generalizable ‘foundation models’ (*47, 48*) evidences that - with enough data - transformers are now broadly seen as the architecture of choice for complex image understanding tasks.

Topography maps were formatted as 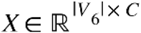 (*V* vertices, *C* cortical spatial maps) and then resampled onto a regularly-tessellated 6th-order icosphere (ico6) with |V_6_| = 40962 vertices and |F_6_| = 40962 patches. This data was ‘tokenised’ by patching the surface with a 2nd-order icosphere |V_2_| = 153 vertices and |F_2_| = 320 non-overlapping triangular patches (Fig. 1). Other icospheric tessellations either decreased model accuracy or did not affect model performance (Supplementary Table 2). This results in an initial input sequence of 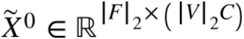. A trainable linear layer compresses each patch to a shorter sequence *(D* = 192), then an extra token of the same dimension was added to the sequence to embed positional encoding, such that the embedded input data was 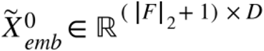. Both reconstruction and translation used a SiT to encode 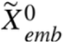:

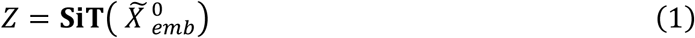

where the **SiT** denotes the multi-head self-attention proposed in (*29*) and adapted to surfaces by (*30*) running multiple self-attention heads (H=3) at each layer (L=6) in parallel. Each head had a dimension of H_d_ = 64. Dropout for positional embeddings and layers were set to 0.3 and 0.5, respectively. As is standard for transformer models, we used LayerNorm (*49*) prior to each multi-head self-attention layer (*29*). Although reconstruction and translation used the same encoder process, they differed in the size of their corresponding linear decoders. Reconstructions (Fig. 1A) needed a bigger linear decoder given the larger size of spatial components:

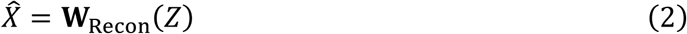

and **W**_*Recon*_(*Z*) ∈ ℝ^*CNV*^. For visualizing reconstruction only, we rearranged outputs to match original brain map dimensions, 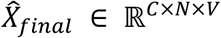 and *C* = 1 given the single spatial map being embedded and reconstructed. For the topography-to-connectome translation models (Fig. 1B), the linear decoders transformed latent embeddings into the upper triangle of connectomes:

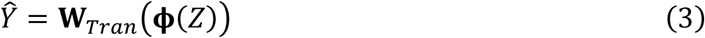

where ϕ (·) is the GELU activation function (*50*). The GELU activation function ensured that translated connectome edges were normally distributed. **W**_*Tran*_ (*Z*) ∈ ℝ^*n*^ and denote the number of vectorized edges along the upper triangle in a connectome. To reconstruct individual differences in spatial topography, all ICA channels were individually normalized by their respective means and standard deviations (calculated from training data) prior to SiT encoding. Concomitantly, the measured ICA topography maps from the validation and test dataset were also normalized using the same training-data means and standard deviations. For translations, only the upper triangle of each subject connectome was used as the ground truth for model optimization. Similarly, all connectomes were demeaned using the training-data means, which were also subtracted from validation and test connectomes. FC is largely symmetrical across hemispheres, so only leveraging one hemisphere of the ICA topography maps would reduce computational costs and redundancy. We chose the left hemisphere because there is slightly more lateralized language and motor activity than in the right hemisphere.

### Model Training and Optimization

All reconstruction and translation models were trained with the same training sub-sample (N_train_ = 7104) for 100 epochs. Model evaluation occurred at each epoch by computing MSE and MAE on the validation sub-sample (N_val_ = 920). After training, only the models with the best validation MSE were chosen for testing (Supplementary Fig. S2). Models with the best validation MSE were chosen over MAE due to better reconstruction and translation performance (Supplementary Table 1). For training, all models minimized the squared L2 norm by comparing element-wise the measured target and the model output (reconstructed/ translated):

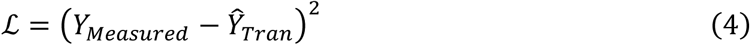

Where Y_Measured_ corresponds to the measured (normalized) vectorized spatial maps for reconstruction, or the vectorized (demeaned) upper triangle of connectomes for translation. We further used a mini-batching size of 32 subjects for faster and more stable convergence. We trained our models with the AdamW optimizer and a cosine decay curve placed on the learning rate (LR = 0.01; LR_min_ = 0.00001). Cosine decay gradually reduces the learning rate following a cosine curve. This allows our model to more carefully fine-tune as it approaches convergence, stabilizing training.

### Identifiability analyses

Two metrics assessed subject identification performance. Average rank percentiles measure the fraction of subjects for which a translated connectome is most similar with its corresponding measured counterpart than with other subjects. First, we calculate translation accuracy by finding the bivariate correlation coefficient between measured and translated connectomes for any given subject *i*, where *k*_*ii*_ = corr(*y*_*i*_, *ŷ*_*i*_). We then repeat this process for the same *y*_*i*_ but against all other comparisons, such that *k*_*ij*_ = corr(*y*_*i*_, *ŷ*_*j*_) and *i* ≠ *j*. Finally, we compute the average rank percentile by:

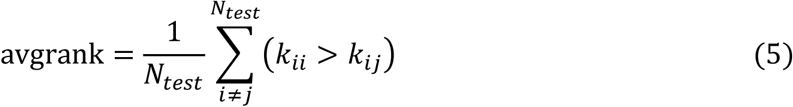

Top-1 Accuracy, instead, is a binary version of average rank percentile. Success only occurs when *k*_*ii*_ > *k*_*ij*_ ∀_*j*_ and *i* ≠ *j*:

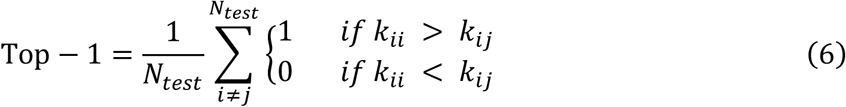

The subject identifiability metrics were repeated for all connectome translations to assess performance for all trained topography-to-connectome models.

### Edge-wise associations between cognition and connectomes

For all connectomes, univariate brain-cognition associations were computed as follows. Across subjects, each edge in the upper triangle of measured and translated connectomes were correlated with cognitive measures, separately. Measured univariate associations and translated univariate associations were then transformed with the Fisher r-to-z transform. Subsequently, the vectorized upper triangle of z-transformed brain-cognition correlations were edge-wise correlated between results from measured connectomes and results from translated connetomes to compute similarity. For both crystallized and fluid intelligence, the aforementioned process was performed separately. All analyses show univariate results for the held-out test sample (N_test_ = 624).

### Multivariate associations for connectomes

We used principal component regressions to reveal multivariate associations between connectome features and cognitive measures. All upper triangle edges were used as connectome features, which are many times bigger than the sample size (N_test_ = 624). Therefore, only the top 100 principal components, calculated within each cross-validation fold, were chosen for dimensionality reduction. In each fold, we then used these 100 components as regressors to train a linear regression model on cognitive scores for crystallized and fluid intelligence, separately. We randomly split the test sample size into a 90/10 split for training and test, respectively, resulting in an R^2^ score for our model. This 90/10 cross validation was repeated 1000 times (‘shuffle splits’), each time producing a respective R^2^ score for measured multivariate associations and translated multivariate associations.

## Acknowledgements

Data used in the preparation of this article were obtained from the Adolescent Brain Cognitive Development (ABCD) Study (https://abcdstudy.org), previously held in the NIMH Data Archive (NDA) and now available from the Lasso data access platform (https://www.nbdc-datahub.org/data-tools-lasso). The ABCD is a multi-site longitudinal study designed to recruit more than 10,000 children ages 9–10 and follow them over 10 years into early adulthood. The ABCD Study is supported by the National Institutes of Health and additional federal partners under award numbers U01DA041022, U01DA041028, U01DA041048, U01DA041089, U01DA041106, U01DA041117, U01DA041120, U01DA041134, U01DA041148, U01DA041156, U01DA041174, U24DA041123, U24DA041147, U01DA041093, and U01DA041025. A full list of supporters is available at https://abcdstudy.org/federal-partners.html. A listing of participating sites and a complete listing of the study investigators can be found at https://abcdstudy.org/scientists/workgroups/. ABCD consortium investigators designed and implemented the study and/or provided data but did not necessarily participate in analysis or writing of this report. This manuscript reflects the views of the authors and may not reflect the opinions or views of the NIH or ABCD consortium investigators. The ABCD data repository grows and changes over time. The ABCD data used in this report came from Annual Release 2.0 (doi: 10.15154/1503209).

## Funding

This work was supported by grants from the National Institutes of Health (NIMH R01 MH128286 & NIMH R01 MH132962; JDB) and the National Science Foundation (Grant No. 2139839; SNR). Computations were performed using the facilities of the Washington University Research Computing and Informatics Facility (RCIF), which has received funding from NIH S10 program grants: 1S10OD025200-01A1 and 1S10OD030477-01.

## Author contributions

Conceptualization: SNR, FA, JB. Methodology: SNR, FA, SS, ER, SD, UK, JB. Investigation: SNR, FA, JB. Visualization: SNR, FA. Data curation: SNR, FA, TE, HM. Supervision: JB. Writing—original draft: SNR, JB. Writing—review & editing: SNR, FA, TE, SS, TG, GK, HM, SD, ER, UK, JB.

## Competing Interests

The authors declare that they have no competing interests.

## Data and materials availability

ABCD data are available at https://www.nbdc-datahub.org/data-tools-lasso. All code used in this article can be found at the public GitHub repo: https://github.com/naranjorincon/ICAsurf2connectomes

## Supplementary Materials

**Figure S1.**
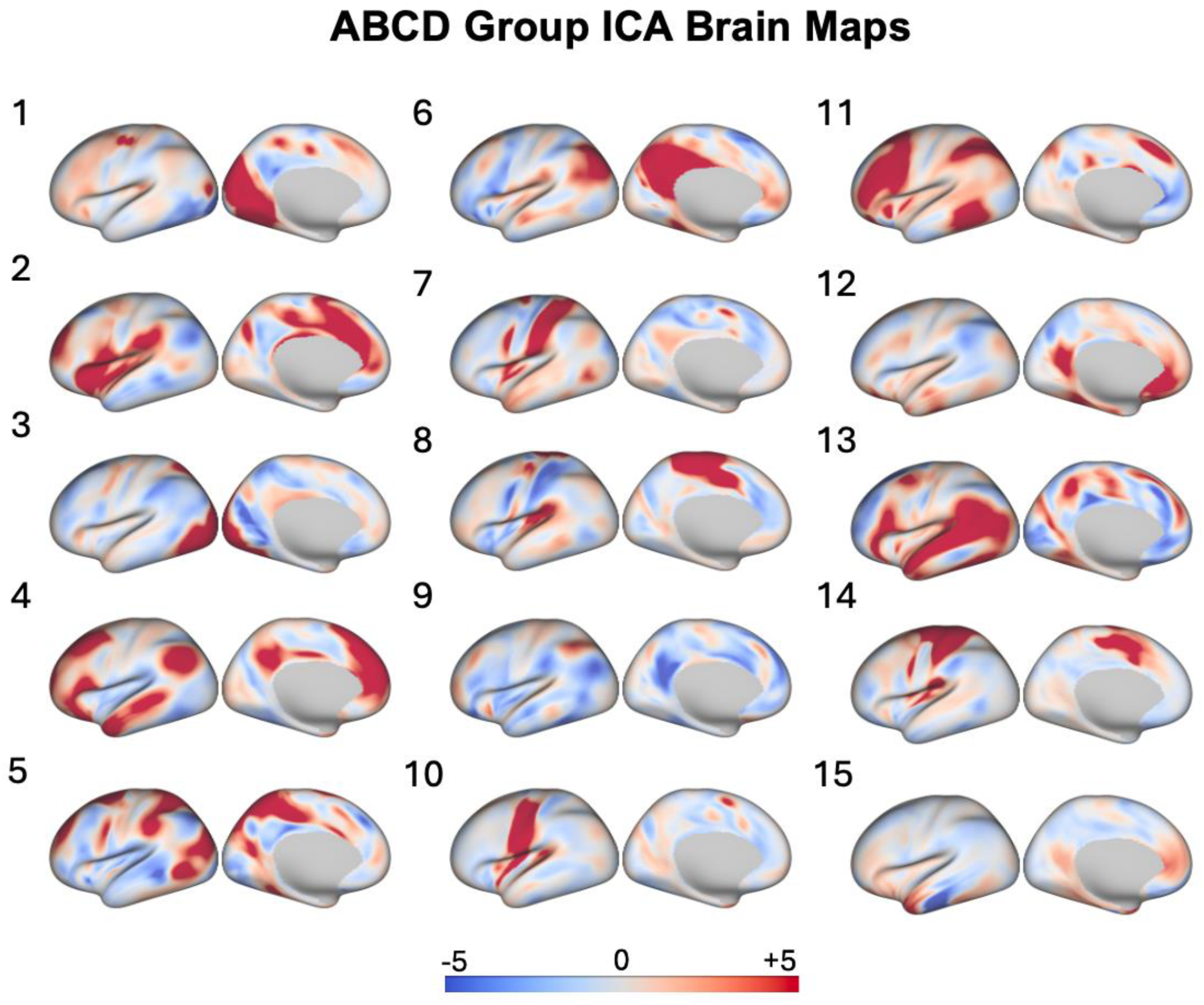
Group ICA maps for ABCD cohort. All ICA brain maps were derived using the FSL toolbox. Melodic was used to decompose rsfMRI scans into 15 independent components.

**Supplementary Table 1.** Topography-to-connectome model selection criteria across translations. To reduce overfitting to the training data, a separate held-out validation sample (N_val_ = 920) was used to compute MSE and MAE. Only models with the lowest MSE and MAE metrics were saved and used for testing. Performance of each selected model on the validation sample is shown on the table for all translations. All models were trained on 10 Intel Xeon Gold 6226R CPUs.

|  | Schaefer-100 | Schaefer-100 PC | Schaefer-300 | Glasser-360 |
| --- | --- | --- | --- | --- |
| MSE | 0.025 ± 0.009 | 0.002 ± 0.001 | 0.028 ± 0.011 | 0.028 ± 0.011 |
| MAE | 0.123 ± 0.021 | 0.033 ± 0.009 | 0.129 ± 0.022 | 0.129 ± 0.022 |

**Figure S2.**
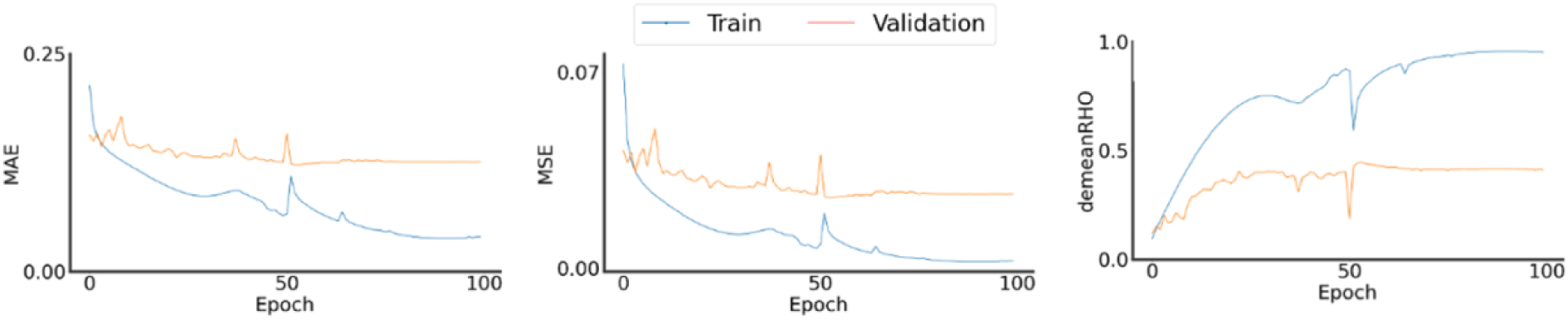
Model Training and Optimization Metrics. Mini-batch training epochs were used to train the topography-to-connectome translation model. MAE and MSE were used to assess performance on a held-out validation dataset. The best MSE model was chosen for all analyses and illustrations. Demeaned correlation results show steady increases as MAE and MSE decrease, but plateau at around the midpoint of training.

**Figure S3.**
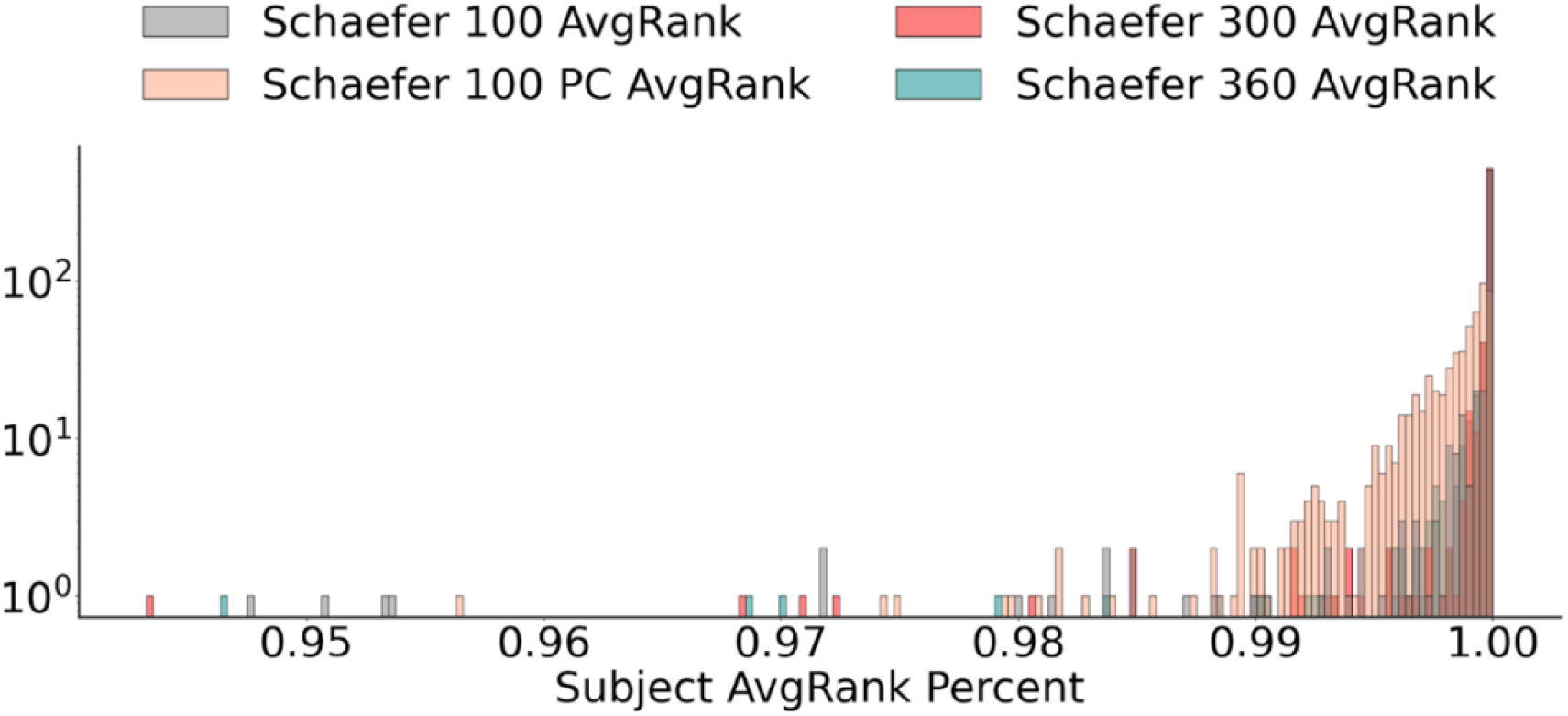
Average rank percentiles for all connectome types. Subject count on a logarithmic scale for average rank percentiles across all connectome types. A percentile of 100% (1.00) is equivalent to perfect subject finger printing based on translated connectomes. All percentiles are based on a held-out dataset of N_test_ = 624.

**Supplementary Table 2.** Topography-to-connectome model performance for Schaefer-100 translations across different ICO resolutions. All models were assessed with MSE and MAE during training. The table shows lowest MSE and MAE metrics achieved by the models during training on the validation sample (N_val_ = 920) across different ICO resolutions. ICO-04 inputs were too computationally demanding, such that a model could not be trained. All models were trained on 10 Intel Xeon Gold 6226R CPUs. OOM = Out Of Memory.

|  | ICO-01 | ICO-02 | ICO-03 | ICO-04 |
| --- | --- | --- | --- | --- |
| MSE | $0.025 \pm 0.01$ | $0.025 \pm 0.01$ | $0.04 \pm 0.02$ | OOM |
| MAE | $0.123 \pm 0.02$ | $0.123 \pm 0.02$ | $0.15 \pm 0.03$ | OOM |
| RHO (demeaned) | $0.43 \pm 0.09$ | $0.45 \pm 0.09$ | $0.32 \pm 0.09$ | OOM |

## Notes

https://github.com/naranjorincon/ICAsurf2connectomes

### Competing Interest Statement

The authors have declared no competing interest.

### Summary of Updates

Corrected affiliations for Emma Robinson and Simon Dahan.

## References

1. O. Sporns, G. Tononi, R. Kötter, The Human Connectome: A Structural Description of the Human Brain. PLOS Comput. Biol. 1, e42 (2005).

2. E. Bullmore, O. Sporns, Complex brain networks: graph theoretical analysis of structural and functional systems. Nat. Rev. Neurosci. 10, 186–198 (2009).

3. O. Sporns, The human connectome: a complex network. Ann. N. Y. Acad. Sci. 1224, 109–125 (2011).

4. A. Griffa, P. S. Baumann, J.-P. Thiran, P. Hagmann, Structural connectomics in brain diseases. NeuroImage 80, 515–526 (2013).

5. M. Cao, H. Huang, Y. He, Developmental Connectomics from Infancy through Early Childhood. Trends Neurosci. 40, 494–506 (2017).

6. A.-L. Goddings, D. Roalf, C. Lebel, C. K. Tamnes, Development of white matter microstructure and executive functions during childhood and adolescence: a review of diffusion MRI studies. Dev. Cogn. Neurosci. 51, 101008 (2021).

7. P. E. Vértes, E. T. Bullmore, Annual Research Review: Growth connectomics – the organization and reorganization of brain networks during normal and abnormal development. J. Child Psychol. Psychiatry 56, 299–320 (2015).

8. P. Hagmann, L. Cammoun, X. Gigandet, R. Meuli, C. J. Honey, V. J. Wedeen, O. Sporns, Mapping the structural core of human cerebral cortex. PLoS Biol. 6, e159 (2008).

9. M. A. Koch, D. G. Norris, M. Hund-Georgiadis, An investigation of functional and anatomical connectivity using magnetic resonance imaging. NeuroImage 16, 241–250 (2002).

10. R. E. Passingham, K. E. Stephan, R. Kötter, The anatomical basis of functional localization in the cortex. Nat. Rev. Neurosci. 3, 606–616 (2002).

11. C. J. Honey, O. Sporns, L. Cammoun, X. Gigandet, J. P. Thiran, R. Meuli, P. Hagmann, Predicting human resting-state functional connectivity from structural connectivity. Proc. Natl. Acad. Sci. 106, 2035–2040 (2009).

12. D. M. Cole, S. M. Smith, C. F. Beckmann, Advances and pitfalls in the analysis and interpretation of resting-state FMRI data. Front. Syst. Neurosci. 4 (2010).

13. D. Wang, R. L. Buckner, M. D. Fox, D. J. Holt, A. J. Holmes, S. Stoecklein, G. Langs, R. Pan, T. Qian, K. Li, J. T. Baker, S. M. Stufflebeam, K. Wang, X. Wang, B. Hong, H. Liu, Parcellating cortical functional networks in individuals. Nat. Neurosci. 18, 1853–1860 (2015).

14. R. Kong, J. Li, C. Orban, M. R. Sabuncu, H. Liu, A. Schaefer, N. Sun, X.-N. Zuo, A. J. Holmes, S. B. Eickhoff, B. T. T. Yeo, Spatial Topography of Individual-Specific Cortical Networks Predicts Human Cognition, Personality, and Emotion. Cereb. Cortex 29, 2533–2551 (2019).

15. J. Bijsterbosch, S. J. Harrison, S. Jbabdi, M. Woolrich, C. Beckmann, S. Smith, E. P. Duff, Challenges and future directions for representations of functional brain organization. Nat. Neurosci. 23, 1484–1495 (2020).

16. X. Li, N. Bianchini Esper, L. Ai, S. Giavasis, H. Jin, E. Feczko, T. Xu, J. Clucas, A. Franco, Sólon Heinsfeld, A. Adebimpe, J. T. Vogelstein, C.-G. Yan, O. Esteban, R. A. Poldrack, C. Craddock, D. Fair, T. Satterthwaite, G. Kiar, M. P. Milham, Moving beyond processing- and analysis-related variation in resting-state functional brain imaging. Nat. Hum. Behav. 8, 2003–2017 (2024).

17. R. Kong, R. N. Spreng, A. Xue, R. F. Betzel, J. R. Cohen, J. S. Damoiseaux, F. De Brigard, S. B. Eickhoff, A. Fornito, C. Gratton, E. M. Gordon, A. J. Holmes, A. R. Laird, L. Larson-Prior, L. D. Nickerson, A. L. Pinho, A. Razi, S. Sadaghiani, J. M. Shine, A. Yendiki, B. T. T. Yeo, L. Q. Uddin, A network correspondence toolbox for quantitative evaluation of novel neuroimaging results. Nat. Commun. 16, 2930 (2025).

18. L. Q. Uddin, B. T. T. Yeo, R. N. Spreng, Towards a Universal Taxonomy of Macro-scale Functional Human Brain Networks. Brain Topogr. 32, 926–942 (2019).

19. R. Kong, Y. R. Tan, N. Wulan, L. Q. R. Ooi, S.-R. Farahibozorg, S. Harrison, J. D. Bijsterbosch, B. C. Bernhardt, S. Eickhoff, B. T. Thomas Yeo, Comparison between gradients and parcellations for functional connectivity prediction of behavior. NeuroImage 273, 120044 (2023).

20. K. Dadi, M. Rahim, A. Abraham, D. Chyzhyk, M. Milham, B. Thirion, G. Varoquaux, Benchmarking functional connectome-based predictive models for resting-state fMRI. NeuroImage 192, 115–134 (2019).

21. J. D. Bijsterbosch, M. W. Woolrich, M. F. Glasser, E. C. Robinson, C. F. Beckmann, D. C. Van Essen, S. J. Harrison, S. M. Smith, The relationship between spatial configuration and functional connectivity of brain regions. eLife 7, e32992 (2018).

22. S. Marek, B. Tervo-Clemmens, F. J. Calabro, D. F. Montez, B. P. Kay, A. S. Hatoum, M. R. Donohue, W. Foran, R. L. Miller, T. J. Hendrickson, S. M. Malone, S. Kandala, E. Feczko, O. Miranda-Dominguez, A. M. Graham, E. A. Earl, A. J. Perrone, M. Cordova, O. Doyle, L. A. Moore, G. M. Conan, J. Uriarte, K. Snider, B. J. Lynch, J. C. Wilgenbusch, T. Pengo, A. Tam, J. Chen, D. J. Newbold, A. Zheng, N. A. Seider, A. N. Van, A. Metoki, R. J. Chauvin, T. O. Laumann, D. J. Greene, S. E. Petersen, H. Garavan, W. K. Thompson, T. E. Nichols, B. T. T. Yeo, D. M. Barch, B. Luna, D. A. Fair, N. U. F. Dosenbach, Reproducible brain-wide association studies require thousands of individuals. Nature 603, 654–660 (2022).

23. T. Easley, K. Freese, E. Munch, J. Bijsterbosch, Using topological data analysis to compare inter-subject variability across resting state functional MRI brain representations. arXiv 2306.13802 [Preprint] (2025). 10.48550/arXiv.2306.13802.

24. J. Dadashkarimi, A. Karbasi, Q. Liang, M. Rosenblatt, S. Noble, M. Foster, R. Rodriguez, B. Adkinson, J. Ye, H. Sun, C. Camp, M. Farruggia, L. Tejavibulya, W. Dai, R. Jiang, A. Pollatou, D. Scheinost, Cross Atlas Remapping via Optimal Transport (CAROT):p Creating connectomes for different atlases when raw data is not available. Med. Image Anal. 88, 102864 (2023).

25. K. W. Jamison, Z. Gu, Q. Wang, C. Tozlu, M. R. Sabuncu, A. Kuceyeski, Krakencoder: a unified brain connectome translation and fusion tool. Nat. Methods 22, 1583–1592 (2025).

26. Y.-F. Tan, J. L. Liow, P.-S. Tan, F. Noman, R. C.-W. Phan, H. Ombao, C.-M. Ting, SFC-GAN: A Generative Adversarial Network for Brain Functional and Structural Connectome Translation. arXiv 2501.07055 [Preprint] (2025). 10.48550/arXiv.2501.07055.

27. L. D. Nickerson, S. M. Smith, D. Öngür, C. F. Beckmann, Using Dual Regression to Investigate Network Shape and Amplitude in Functional Connectivity Analyses. Front. Neurosci. 11 (2017).

28. C. F. Beckmann, S. M. Smith, Probabilistic independent component analysis for functional magnetic resonance imaging. IEEE Trans. Med. Imaging 23, 137–152 (2004).

29. A. Vaswani, N. Shazeer, N. Parmar, J. Uszkoreit, L. Jones, A. N. Gomez, L. Kaiser, I. Polosukhin, Attention Is All You Need. arXiv 1706.03762 [Preprint] (2023). 10.48550/arXiv.1706.03762.

30. S. Dahan, A. Fawaz, L. Z. J. Williams, C. Yang, T. S. Coalson, M. F. Glasser, A. D. Edwards, D. Rueckert, E. C. Robinson, Surface Vision Transformers: Attention-Based Modelling applied to Cortical Analysis. arXiv 2203.16414 [Preprint] (2022). 10.48550/arXiv.2203.16414.

31. A. Schaefer, R. Kong, E. M. Gordon, T. O. Laumann, X.-N. Zuo, A. J. Holmes, S. B. Eickhoff, B. T. T. Yeo, Local-Global Parcellation of the Human Cerebral Cortex from Intrinsic Functional Connectivity MRI. Cereb. Cortex 28, 3095–3114 (2018).

32. E. C. Robinson, S. Jbabdi, M. F. Glasser, J. Andersson, G. C. Burgess, M. P. Harms, S. M. Smith, D. C. Van Essen, M. Jenkinson, MSM: A new flexible framework for Multimodal Surface Matching. NeuroImage 100, 414–426 (2014).

33. T. S. Coalson, D. C. Van Essen, M. F. Glasser, The impact of traditional neuroimaging methods on the spatial localization of cortical areas. Proc. Natl. Acad. Sci. 115, E6356–E6365 (2018).

34. J. Dafflon, P. F. Da Costa, F. Váša, R. P. Monti, D. Bzdok, P. J. Hellyer, F. Turkheimer, J. Smallwood, E. Jones, R. Leech, A guided multiverse study of neuroimaging analyses. Nat. Commun. 13, 3758 (2022).

35. E. S. Finn, X. Shen, D. Scheinost, M. D. Rosenberg, J. Huang, M. M. Chun, X. Papademetris, R. T. Constable, Functional connectome fingerprinting: identifying individuals using patterns of brain connectivity. Nat. Neurosci. 18, 1664–1671 (2015).

36. M. G. Puxeddu, M. Pope, T. F. Varley, J. Faskowitz, O. Sporns, Leveraging multivariate information for community detection in functional brain networks. Commun. Biol. 8, 840 (2025).

37. G. Marrelec, A. Krainik, H. Duffau, M. Pélégrini-Issac, S. Lehéricy, J. Doyon, H. Benali, Partial correlation for functional brain interactivity investigation in functional MRI. NeuroImage 32, 228–237 (2006).

38. M. Fiecas, H. Ombao, D. van Lunen, R. Baumgartner, A. Coimbra, D. Feng, Quantifying temporal correlations: A test–retest evaluation of functional connectivity in resting-state fMRI. NeuroImage 65, 231–241 (2013).

39. K. L. Peterson, R. Sanchez-Romero, R. D. Mill, M. W. Cole, Regularized partial correlation provides reliable functional connectivity estimates while correcting for widespread confounding. Imaging Neurosci. 3, IMAG.a.162 (2025).

40. B. J. Casey, T. Cannonier, M. I. Conley, A. O. Cohen, D. M. Barch, M. M. Heitzeg, M. E. Soules, T. Teslovich, D. V. Dellarco, H. Garavan, C. A. Orr, T. D. Wager, M. T. Banich, N. K. Speer, M. T. Sutherland, M. C. Riedel, A. S. Dick, J. M. Bjork, K. M. Thomas, B. Chaarani, M. H. Mejia, D. J. Hagler, M. Daniela Cornejo, C. S. Sicat, M. P. Harms, N. U. F. Dosenbach, M. Rosenberg, E. Earl, H. Bartsch, R. Watts, J. R. Polimeni, J. M. Kuperman, D. A. Fair, A. M. Dale, The Adolescent Brain Cognitive Development (ABCD) study: Imaging acquisition across 21 sites. Dev. Cogn. Neurosci. 32, 43–54 (2018).

41. E. Feczko, G. Conan, S. Marek, B. Tervo-Clemmens, M. Cordova, O. Doyle, E. Earl, A. Perrone, D. Sturgeon, R. Klein, G. Harman, D. Kilamovich, R. Hermosillo, O. Miranda-Dominguez, A. Adebimpe, M. Bertolero, M. Cieslak, S. Covitz, T. Hendrickson, A. C. Juliano, K. Snider, L. A. Moore, J. Uriartel, A. M. Graham, F. Calabro, M. D. Rosenberg, K. M. Rapuano, B. J. Casey, R. Watts, D. Hagler, W. K. Thompson, T. E. Nichols, E. Hoffman, B. Luna, H. Garavan, T. D. Satterthwaite, S. F. Ewing, B. Nagel, N. U. F. Dosenbach, D. A. Fair, Adolescent Brain Cognitive Development (ABCD) Community MRI Collection and Utilities. bioRxiv [Preprint] (2021). 10.1101/2021.07.09.451638.

42. M. Goncalves, J. Moser, T. J. Madison, R. McCollum, J. T. Lundquist, B. Fayzullobekova, L. Hadera, H. H. N. Pham, L. A. Moore, A. Houghton, G. Conan, M. A. Styner, D. Alexopoulos, C. D. Smyser, S. M. Stoyell, S. Koirala, S. M. Nelson, K. B. Weldon, E. Lee, R. J. M. Hermosillo, L. Vizioli, E. Yacoub, G. H. Patel, J. Sanchez, K. Wengler, T. Salo, T. D. Satterthwaite, J. T. Elison, C. J. Markiewicz, R. A. Poldrack, E. Feczko, O. Esteban, D. A. Fair, fMRIPrep Lifespan: Extending A Robust Pipeline for Functional MRI Preprocessing to Developmental Neuroimaging. bioRxiv [Preprint] (2025). 10.1101/2025.05.14.654069.

43. M. F. Glasser, T. S. Coalson, E. C. Robinson, C. D. Hacker, J. Harwell, E. Yacoub, K. Ugurbil, J. Andersson, C. F. Beckmann, M. Jenkinson, S. M. Smith, D. C. Van Essen, A multi-modal parcellation of human cerebral cortex. Nature 536, 171–178 (2016).

44. S. Dahan, L. Z. J. Williams, D. Rueckert, E. C. Robinson, The Multiscale Surface Vision Transformer. arXiv 2303.11909 [Preprint] (2024). 10.48550/arXiv.2303.11909.

45. A. Dosovitskiy, L. Beyer, A. Kolesnikov, D. Weissenborn, X. Zhai, T. Unterthiner, M. Dehghani, M. Minderer, G. Heigold, S. Gelly, J. Uszkoreit, N. Houlsby, An Image is Worth 16x16 Words: Transformers for Image Recognition at Scale. arXiv 2010.11929 [Preprint] (2021). 10.48550/arXiv.2010.11929.

46. A. Fawaz, L. Z. J. Williams, A. Alansary, C. Bass, K. Gopinath, M. da Silva, S. Dahan, C. Adamson, B. Alexander, D. Thompson, G. Ball, C. Desrosiers, H. Lombaert, D. Rueckert, A. D. Edwards, E. C. Robinson, Benchmarking Geometric Deep Learning for Cortical Segmentation and Neurodevelopmental Phenotype Prediction. bioRxiv [Preprint] (2021). 10.1101/2021.12.01.470730.

47. J. L. Ba, J. R. Kiros, G. E. Hinton, Layer Normalization. arXiv 1607.06450 [Preprint] (2016). 10.48550/arXiv.1607.06450.

48. D. Hendrycks, K. Gimpel, Gaussian Error Linear Units (GELUs). arXiv 1606.08415 [Preprint] (2023). 10.48550/arXiv.1606.08415.

